# Incubation of craving for alcohol-associated cues is reduced by running-wheel exercise

**DOI:** 10.64898/2026.04.25.720858

**Authors:** Thomas M Ferella, Alexander IJ Kilby, Laisa de Siquiera Umpierrez, Amy O’Connor, Mia Swinberg, Andrew J Lawrence, Jennifer L. Cornish, Christina J Perry

**Affiliations:** School of Psychological Sciences, Macquarie University, North Ryde, NSW 2113, Australia; The Florey Institute of Neuroscience and Mental Health, Parkville, Melbourne, VIC 3052 Australia; Florey Department of Neuroscience and Mental Health, The University of Melbourne, Parkville, VIC 3052, Australia

**Keywords:** Alcohol, drug-seeking, cue-induced reinstatement, exercise, reward circuit

## Abstract

Craving – the powerful urge to seek and consume alcohol in response to alcohol-associated cues does not diminish after drinking cessation but rather is magnified throughout abstinence. This phenomenon, termed “incubation of craving”, contributes to the relapsing nature of alcohol use disorder. Despite its occurrence in human populations and being well-studied in rodent models of psychostimulant drug relapse, the underlying neural mechanisms and potential treatments remain largely unexplored for alcohol-related incubation of craving. Our research seeks to meet this gap, and this particular study investigated the neural correlates of the incubation of craving for alcohol-associated cues and assessed whether exercise could prevent increased relapse propensity in rats.

Male Long Evans rats were trained to lever press for an alcohol reward delivered with simultaneous presentation of a discrete cue. This response was then extinguished and reinstated by presenting the discrete cue alone when rats pressed the lever. Cue-induced reinstatement occurred either on day 1 following extinction (No Abstinence) or on day 29 (Abstinence). A third group was tested on day 29 and had 4-hour daily voluntary running wheel access throughout this abstinence period (Exercise). All rats were perfused 90 minutes following test, and relative activation across the brain was estimated by quantifying c-Fos protein immunoreactivity. The brain-wide coordination of neural activity was also mapped.

We found a robust incubation of craving effect for alcohol-associated cues, which was mitigated by exercise. Immunohistochemistry revealed that the Abstinence group demonstrated higher c-Fos immunoreactivity compared to the No Abstinence group in multiple reinstatement-related brain regions. This effect was reversed in the Exercise group. Brain-wide neural mapping demonstrated that the Abstinence group had decreased modularity (groups of coordinating brain regions) compared to the no-abstinence group. Although network connectivity profile in the exercise group was different from no abstinence, we found that overall neural activation returned to a similar modularity profile of clustered regions as this condition, indicating that exercise does not attenuate the incubation of craving effect by reversing all the neural effects of abstinence. Rather, exercise may be acting upon select brain regions or pathways to exert relapse protective effects by restoring widespread interconnectivity. This is the first study to investigate neural activation in incubated alcohol-seeking, and provides supporting evidence for promoting voluntary exercise as an adjunctive treatment for relapse prevention in alcohol-use disorder.

---

Relapse is a cardinal feature of alcohol use disorder, and is usually preceded by a powerful urge to seek and consume alcohol: “craving” (Sliedrecht et al., 2019). Alcohol-associated cues can trigger craving, even after extended periods without alcohol consumption, something that is common to all substance use disorders. In fact, the physiological response to drug-associated cues is reported to increase rather than fade with increasing lengths of abstinence. This phenomenon, termed ‘incubation of craving’ (Gawin & Kleber, 1986) has been reported across multiple drugs of abuse, including nicotine (Bedi et al., 2011), methamphetamine (Wang et al., 2013; Zhao et al., 2021) and alcohol (Bach et al., 2020; Li et al., 2015; Treloar Padovano & Miranda, 2021), and is a challenge for enduring remission (Parvaz et al., 2016; Pickens et al., 2011; Zhao et al., 2021).

Given that cue-induced drug seeking has long been considered an operational measure of craving in rodents and other preclinical models (Markou et al., 1993), incubation of craving is modelled through the time-dependent increases in cue-elicited drug-seeking responses following periods of abstinence (Grimm et al., 2001). In rodent models, this effect is observed with cues previously paired with the self-administration of cocaine (Grimm et al., 2001; Lu et al., 2004; Ma et al., 2014), nicotine (Funk et al., 2016; Markou et al., 2018), methamphetamine (Adhikary et al., 2017; Shepard et al., 2004), and opiates (Altshuler et al., 2021; Patel & Loweth, 2024; Shalev et al., 2001).

While many studies have investigated cue-induced alcohol relapse and the implicated neurocircuitry (Keistler et al., 2017; Koob & Volkow, 2016; Perry et al., 2014), very few studies employ models to compare neural changes underlying cue-induced reinstatement responding pre- and post-abstinence (i.e. incubation). A review of the literature uncovered only two preclinical rodent studies that demonstrably investigated the incubation of craving in the context of alcohol seeking and relapse. In outbred rats trained to lever press for alcohol, cue-induced responding to the alcohol-associated cue was greater on abstinence day 56 compared to abstinence day 1 (Bienkowski et al., 2004). On the other hand, Jupp et al. (2011) found no behavioural differences during cue-induced reinstatement of alcohol-seeking after 5 months of abstinence compared to no abstinence. In the latter, despite the lack of behavioural differences, neural activation of the medial prefrontal, piriform and orbitofrontal cortices was increased during the delayed compared to the immediate test (Jupp et al., 2011). This is similar to the increased recruitment of reward circuitry observed for incubation for heroin (Fanous et al., 2012), oxycodone (Altshuler et al., 2021), methamphetamine (Caprioli et al., 2017; Davis et al., 2021; Li et al., 2018; Rossi et al., 2020), and nicotine (Funk et al., 2016) craving. However, to date, the underlying neural mechanisms of incubation of craving for alcohol remain largely uncharacterised.

Mouse models utilising chronic intermittent ethanol vapour exposure have revealed that withdrawal and abstinence from alcohol causes brain wide changes in regional c-Fos recruitment and connectivity (Kimbrough et al., 2020; Roland et al., 2023; Smith et al., 2020). Using hierarchical clustering analyses, these studies identified a reduction in the number of independent clusters of brain regions coordinating together following withdrawal, that is, neural activity of different brain regions was more related. This suggests that alcohol withdrawal leads to enhanced coordination of brain wide circuitry and provides strong evidence of withdrawal-induced plasticity. However, in all of these studies, alcohol exposure was via passive and enforced vapour inhalation. To fully understand how abstinence affects craving, it is important to also look at similar functional connectivity changes in the context of voluntary alcohol-seeking behaviour.

The prevalence of incubation in clinical populations (Bach et al., 2020; Li et al., 2015; Treloar Padovano & Miranda, 2021), also highlights the need for preventative treatments to reduce relapse rates. In rodent self-administration models, exercise (via voluntary wheel-running access) has been demonstrated to reduce cue elicited drug-seeking for cocaine, methamphetamine, and heroin (Lynch et al., 2010; Sanchez et al., 2013; Smethells et al., 2020; Zlebnik et al., 2010). Furthermore, exercise attenuated the incubation of craving at 30 vs 3 days of abstinence from cocaine in both male and female rats (Carroll et al., 2022; Zlebnik & Carroll, 2015). There also is substantial evidence that voluntary exercise reduces alcohol intake and preference in both male and female rodents (Booher et al., 2019; Brager & Hammer, 2012; Darlington et al., 2014; Darlington et al., 2016; Ehringer et al., 2009; Gallego et al., 2015; Pichard et al., 2009). Furthermore, combined cue- and context-induced reinstatement of alcohol seeking, tested 4 weeks from the final day of self-administration, was attenuated if rats had access to running wheels throughout this time (Somkuwar et al., 2016). It’s worth noting however, that in this study cued alcohol-seeking responses were not measured at the beginning of abstinence, hence any effect of wheel running on incubation – the increase in such responses across abstinence – cannot be inferred per se. Indeed, to our current knowledge there remains no published findings regarding exercise as a treatment specifically for the incubation of craving for alcohol-associated cues.

The present research aimed to assess the changes in neural activation and coordination associated with the time-dependent increases in responding to alcohol-associated cues across the brain, and how this is affected by exercise throughout abstinence.

## General methods

### Animals and Housing

Male Long Evans rats aged six to eight weeks (N = 24) were ordered from the Animal Resources Centre (Perth, WA). Rats were housed in groups of 4 in standard mesh wire-top cages (64cm x 40cm x 20cm). All animals were housed in temperature (21°C ± 1°C) and humidity-controlled (60%) rooms under a 12-hr reverse light cycle (lights on 07:00pm). All cages contained corn cob and shredded paper bedding, perspex tunnels, and wood blocks for enrichment. Rats were provided food and water ad libitum except during operant training procedures Upon arrival, all animals underwent habituation to their environment and were handled daily for 1 week prior to commencement of experimental procedures.

All procedures conducted for this experiment were approved by the Macquarie University Animal Ethics Committee (ARA 2020-034) and followed the guidelines of the National Health and Medical Research Council Australian Code for the Care and Use of Animals for Scientific Purposes (2013), and in accordance with the Prevention of Cruelty to Animals Act (2004).

### Apparatus

Self-administration, extinction and cue-induced reinstatement procedures occurred in standard operant conditioning chambers (30 x 24 x 29cm; Med Associates, VT, USA) as previously described (Robinson et al., 2022; Umpierrez et al., 2026), with the addition of a singular liquid magazine catchment between levers for solution delivery. Abstinence procedures were conducted in single-housed, plexiglass chambers (Techniplast, Buguggiate, VA, Italy; 22 x 43 x 20cm), with mesh wire tops containing either an inbuilt running wheel (106cm circumference) or no wheel. A magnetic ticker was used to record the number of wheel rotations. Food and water were provided *ad libitum* in both chambers.

### Experimental Design

#### Self-Administration, Extinction, and Reinstatement Procedures

Self-administration sessions were conducted 5 days a week (Monday-Friday), while extinction sessions were conducted daily; all sessions 30 minutes in length. Left and right levers were assigned as ‘active’ or ‘inactive’ (counterbalanced). During self-administration, presses on the active lever activated an external infusion pump resulting in delivery of ∼170 μl of a sweetened alcohol solution (6% v/v alcohol, 5% w/v maltodextrin in water) into the magazine. Simultaneously, the house light was switched off and the cue light above the active lever was illuminated for 8s to indicate the availability of the alcohol reinforcer. Following each reinforced lever press there was a 20 second timeout during which any further responses to the active lever had no programmed consequences and the house light remained off.

Rats were initially trained to lever press for alcohol under a fixed-ratio (FR)-1 schedule of reinforcement, whereby every active lever press outside of the timeout period was reinforced. This continued for a minimum of 15 days and until acquisition criterion was met (≥ 8 infusions in a single session), before moving to a FR-3 schedule (every third active lever press reinforced) for 5 days. Six rats failed to reach criterion for acquisition and were excluded.

Following self-administration, all rats underwent 7 consecutive daily extinction sessions. During these sessions, the house light was switched off and responses to the active and inactive lever had no programmed consequences.

Subsequently, rats (N = 18) were quasi-randomly allocated to one of three groups: No Abstinence, Abstinence, or Exercise (n = 6/group), such that the mean and standard deviation of lever presses for the last day of self-administration and the first and last day of extinction were closely balanced between groups. Rats underwent a cue-induced reinstatement test either one (No Abstinence) or 29 days (Abstinence and Exercise) following the final extinction session. These sessions were identical to a FR-1 self-administration session, except that the primary alcohol reinforcement solution was not delivered upon lever press. As such, active lever presses led only to the illumination of the cue light on a FR-1 + timeout schedule.

#### Abstinence/Exercise Procedures

Groups Abstinence and Exercise were both subjected to 28 days of forced abstinence following extinction. During this time, rats from group Exercise were placed into the abstinence chambers equipped with the inbuilt running wheel, which were absent for group Abstinence, for 4 hours per day, 7 days a week. A timeline of all behavioural procedures is depicted in Figure 1A.

**Figure 1:**
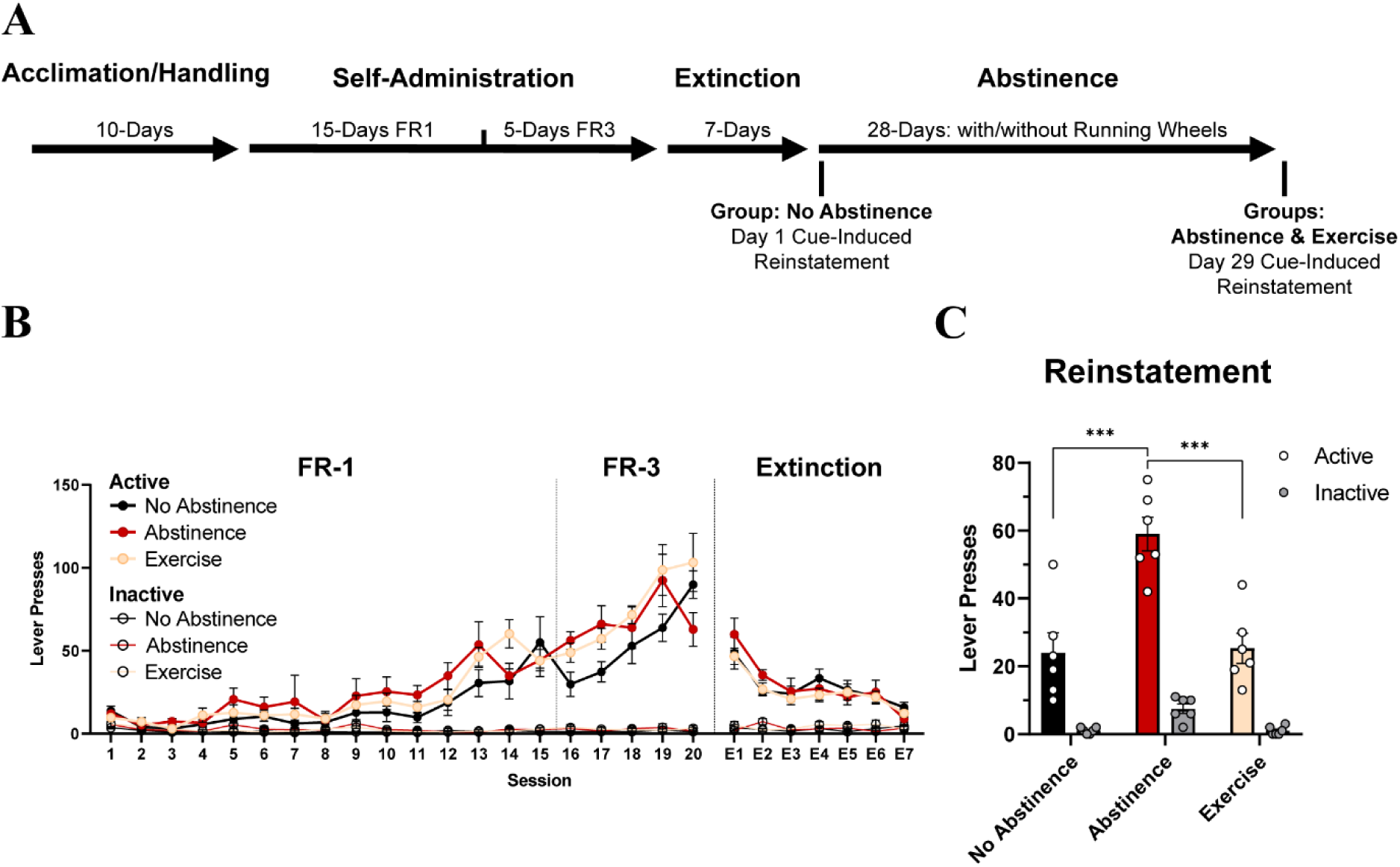
Behaviour. (A) Timeline of behavioural procedures. (B) Lever presses across FR-1, FR-3, and extinction training. No differences in responding were found between the to-be allocated groups across training sessions. (C) Lever presses in 30-min cue-induced reinstatement session for each abstinence group. All data are represented as mean (± SEM). *** *p* < 0.001.

#### Sacrifice and Immunohistochemistry

Neuronal activation during reinstatement sessions was estimated by examining relative expression of c-Fos protein, which is maximally expressed 90 – 120 minutes after cell activation (Kovacs, 2008). Therefore, ninety minutes after the onset of the respective cue-induced reinstatement test session, all rats were deeply anaesthetized via i.p. injections of sodium pentobarbital (100 mg/kg; Lethabarb) and perfused transcardially with 100 mL heparinised saline (10 IU) followed by 250 ml of 4% paraformaldehyde (PFA) w/v in phosphate buffered saline (PBS). Brains were extracted, post-fixed in PFA (24 hours) and then transferred into cryoprotectant (30% ethylene glycol, 30% sucrose, 2% polyvinylpyrrolidone in PBS) and stored at -20°C before being sectioned coronally at 40μm into four adjacent series using a Vibratome (Leica Microsystems, VIC, Australia). Sections were stored again in cryoprotectant at -20°C until immunohistochemistry procedures began.

Chromogenic immunohistochemistry procedures were conducted on one of the four series for each brain, as described previously (Park et al., 2020), utilising a monoclonal c-Fos primary antibody raised in rabbit (1:8000; Cell Signalling Technology, US, Cat# 5348S) and biotinylated goat anti-rabbit secondary (1:500; Vector Laboratories, US, Cat# BA-1000). Sections were mounted onto positively charged microscope slides, air dried, dehydrated and cover slipped. Brightfield photomicrographs of each section were captured using the 10x objective of a Zeiss Axio Imager (Zeiss, Germany), or VS200 Research Slide Scanner (Olympus Life Science, Japan).

#### Quantification of c-Fos immunoreactivity

c-Fos immunoreactivity was quantified bilaterally using the QUINT workflow (Yates et al., 2019) in coronal sections ranging from Bregma +4.68mm To Bregma -4.20mm. Sections were first registered in 3D space to the Waxholm Space atlas of the rat brain version 4.0 (WHS atlas; (Kleven et al., 2023)) accounting for the angle of sectioning (using QuickNII version 2.2 software), and tissue distortion (using VisuAlign version 0.9; (Puchades et al., 2019)). Cell markers of interest were identified using machine learning (Ilastik version 1.4.0; (Berg et al., 2019)). Finally, the output from these three processes was combined to automatically quantify c-Fos expression in all brain regions (Nutil v0.8.0; (Groeneboom et al., 2020)). Full procedure details are available in Supplementary information: QUINT Workflow & Figure S1.

#### Data Analysis

All statistical analyses were conducted using Stata, version 18.0 (StataCorp, College Station, TX, USA) and graphs were produced using GraphPad Prism, version 9.4.1 (GraphPad software Inc., CA, USA), except where explicitly stated otherwise. For all tests, the overall Type 1 error rate was set at 0.05.

#### Behavioural Analysis

Active and inactive lever presses were analysed via separate 3-way mixed-design ANOVAs for acquisition, extinction, and reinstatement. For multi-level ‘Day’ factors, planned orthogonal linear polynomial contrast weights were tested. For the three-level ‘Group’ factor, we tested a set of planned pairwise comparisons that preserved the factorial design, and family-wise error rate was controlled using Bonferroni’s correction. Assumptions of normality were assessed using Shapiro-Wilk tests, homogeneity of variance was assessed using Levene’s test, and Lawley’s test was used to assess compound symmetry to infer sphericity. Assumptions were checked and satisfied prior to each analysis. Normality was violated in some measures; however, the ANOVA model was retained in all instances as group sizes were equal. Where the assumption of sphericity was violated, a Greenhouse-Geisser correction was applied.

#### Neural Analysis

Due to challenges associated with generating a three-dimensional atlas using diffusion tensor and magnetic resonance imaging techniques (Kleven et al., 2023), some brain region divisions are divergent from those in the more commonly used atlas (Paxinos & Watson, 2007). Namely, some common delineations are grouped (e.g. the septal region (Sep) including all medial and lateral septum divisions, and the caudate putamen (CPu) including the dorsomedial- and dorsolateral striatum), while others remain ‘unspecified’. For example, the amygdaloid area unspecified (Am-u) that encompasses all the traditional subdivisions of the amygdala; the basal forebrain region unspecified (BFR-u) which maps on to the diagonal band, surrounds of the ventral pallidum, and extends caudally into the preoptic hypothalamus; the ventral striatal region unspecified (VSR-u) consisting of the lateral nucleus accumbens shell and interstitial nucleus of the posterior limb of the anterior commissure (IPAC); and the hypothalamic region unspecified (HTh-u) which includes the paraventricular and medial hypothalamic nuclei. Supplementary Table S1 outlines a full list of the included WHS atlas regions and their abbreviations.

For each region, c-Fos density was calculated by dividing the total number of immunoreactive cells by the sum of area of the region (mm^2^) across all sections quantified. For regions that form the canonical reward-seeking circuit (Koob & Volkow, 2016; Namba et al., 2018), difference between abstinence conditions was determined via separate one-way between-group ANOVAs, with Tukey’s HSD applied to post-hoc comparisons.

Functional connectivity was explored by conducting hierarchical clustering analyses to visualise modules of brain regions with similar patterns of coordinated activity based on the Euclidean distances of Pearson’s correlations of c-Fos density between each brain region (G. Anversa et al., 2023; Kimbrough et al., 2020; Richards et al., 2025). Prior to analyses, any brain regions that were missing data across all three groups, were unquantifiable for ≥ 4 animals within each group due to damage/loss in histological procedures, as well as all white matter tracts, were excluded across all groups. Additionally, select subregions were combined to be represented as one region based on the WHS atlas divisions (Kleven et al., 2023). Following exclusions and subregion groupings, the final sample consisted of 227 sections for the No Abstinence group (n = 6, range: 36-42 sections per brain), 238 sections for the Abstinence group (n = 6, range: 35-48), and 241 sections for the Exercise group (n = 6, range: 36-46), for a total of 706 sections. A total of 72 brain regions were retained for analyses for each abstinence condition.

For each group, No Abstinence, Abstinence, and Exercise, inter-regional correlations of c-Fos densities were calculated using Pearson’s correlation analysis in GraphPad Prism, version 9.4.1. Euclidean distances for each correlation were computed in SPSS version 29.0.0.0 (IBM, NY, USA). Heatmaps were generated, whereby regions were organised based on the dissimilarity of Euclidean distances using the complete linking method to identify modules of coordination, and clusters were trimmed at 50% dendrogram height. Hierarchical clustering analyses and heatmap generation were performed using Python (version 3.11.5) in JupyterLab (version 3.6.3).

To examine abstinence- and exercise-associated changes in functional connectivity, we adopted a graph theory-based approach to assess the relative contribution of key brain regions driving inter- and intra-modular connectivity in the modules identified in the hierarchical clustering analyses, similarly described in (Kimbrough et al., 2020). We adopted two widely used centrality metrics for modular systems: the within module degree z-score (WMDz) and participation coefficient (PC), as originally defined by Guimera and Nunes Amaral (2005). Graph visualisation was conducted using Python (version 3.11.5) in JupyterLab (version 3.6.3).

Prior to calculating the WMDz and PC metrics, we took the absolute values of Pearson correlations, and applied a threshold of *r* > 0.65, such that every node (brain region) had at least 1 edge (correlation) with another node. Absolute values of the edges were retained for analyses as we sought to determine the strength, rather than directionality per se, of functional connectivity. In brief, the WMDz represents the normalised strength of a given brain region’s connection to its own module (higher WMDz), or other modules (lower WMDz); and the PC represents a measure of inter-module connectivity, where a value of 0 represents that the brain region has connections only to other regions in its own module, and value of 1 if its connections are uniformly distributed across all modules. Values > 0.75 were considered strong WMDz scores, and values greater than 0.5 were considered strong PC scores. See Supplemental Graph Theory Metrics for full details.

## Results

### Behaviour

#### Self-Administration

Mean daily lever presses across self-administration, extinction and reinstatement tests are shown in Figure 1B & 1C. Self-administration data were analysed using a 20 × 2 × 3 (Day × Lever Type × Group) mixed design ANOVA, with Day and Lever Type as within-subjects variables and Group as the between-subjects variable for the self-administration data. Across acquisition, rats increased responding on the active lever only, confirmed by significant main effects for both Lever Type (*F*_[1, 15]_ = 261.86, *p* < 0.001, *η_p_^2^*= 0.946) and Day (*F*_[19, 278]_ = 35.56, *p* < 0.001, *η_p_^2^*= 0.708) and a significant interaction between Day and Lever Type (*F*_[19, 278]_ = 40.00, *p* < 0.001, *η_p_^2^*= 0.732). There were no systematic differences between groups prior to abstinence group allocation, which was confirmed by non-significant Group main effects (*F*_[2, 15]_ = 2.27, *p* = 0.138, *η_p_^2^*= 0.232). Likewise, all interactions with Group were not significant (Lever Type by Group (*F*_[2, 15]_ = 2.02, *p* = 0.167, *η_p_^2^* = 0.212), Day by Group (*F* = 1.52, *p* = 0.136, *η_p_^2^* = 0.172), Day by Lever Type by Group (*F*_[38, 278]_ = 1.66, *p* = 0.093, *η_p_^2^*= 0.185)) (Figure 1B).

#### Extinction

Extinction data were analysed similarly, using a 7 × 2 × 3 (Day × Lever Type × Group). Responding on the active lever decreased across extinction days, confirmed by significant main effects for Lever Type (*F*_[1, 15]_ = 180.33, *p* < 0.001, *η_p_^2^* = 0.923) and Day (*F*_[6, 88]_ = 17.83, *p* < 0.001, *η_p_^2^* = 0.549), and interaction between Day and Lever Type (*F*_[6, 88]_ = 18.35, *p* < 0.001, *η_p_^2^* = 0.556). Again, all Group effects were non-significant (main effect (*F*_[2, 15]_ = 0.21, *p* = 0.812, *η_p_^2^* = 0.027), Lever Type by Group (*F*_[2, 15]_ = 0.66, *p* = 0.530, *η_p_^2^*= 0.081), Day by Group (*F*_[12, 88]_ = 0.82, *p* = 0.590, *η_p_^2^*= 0.100) and Day by Lever Type by Group (*F*_[12, 88]_ = 0.83, *p* = 0.577, *η_p_^2^* = 0.102) (Figure 1B).

#### Exercise

For the Exercise group, the number of wheel rotations was converted to distance run in metres (m). Distance run increased over the abstinence period, from an average (± SEM) of 1639.5 (± 230.9) m/day in week one to 4641.7 (± 176.7) m/day in week 4 (see supplementary Figure S2). Linear regression analyses revealed a non-significant negative association between mean distance run/day and active lever responses in cue-induced reinstatement (*β* = −0.006, *t*_[4]_ = −1.60, *p* = .185, *R^2^* = 0.39).

#### Cue-Induced Reinstatement

Reinstatement data were analysed using a 2 x 2 x 3 (Test Day × Lever Type × Group) mixed model ANOVA, with Test Day and Lever Type as within-subjects variables and Group as the between-subjects variable. Overall, rats responded more to the active lever at cue-induced reinstatement test than on the final day of extinction (Test Day (*F*_[1, 15]_ = 98.89, *p* < 0.001, *η_p_^2^* = 0.868), Lever Type (*F*_[1, 15]_ = 129.89, *p* < 0.001, *η_p_^2^* = 0.896), Test Day by Lever Type (*F*_[1, 15]_ = 82.78, *p* < 0.001, *η_p_^2^* = 0.847)). Reinstatement was different depending on Group, confirmed by a significant main effect for Group (*F*_[2, 15]_ = 12.13, *p* < 0.001, *η_p_^2^* = 0.618), as well as Lever Type by Group (*F*_[2, 15]_ = 5.81, *p* = 0.014, *η_p_^2^* = 0.436), Test Day by Group (*F*_[2, 15]_ = 26.02, *p* < 0.001, *η_p_^2^* = 0.776) and Test Day by Lever Type by Group (*F*_[2, 15]_ = 12.18, *p* < 0.001, *η_p_^2^* = 0.619) interactions. Planned comparisons between each abstinence group revealed that this interaction was driven by greater active responding in reinstatement for the Abstinence compared to the No Abstinence group (*F*_[1, 15]_ = 19.58, *p* < 0.001) and for Abstinence compared to Exercise (*F*_[1, 15]_ = 16.84, *p* < 0.001), while no significant difference between No Abstinence and Exercise groups (*F*_[1, 15]_ = 0.10, *p* = 0.753) was found. Thus, alcohol-seeking increased following four weeks of abstinence, but this effect was absent where rats were given access to running wheels in the intervening period.

#### Neural Findings

##### c-Fos expression

Across the selected ROIs canonically implicated in reward-seeking behaviour, analyses showed that c-Fos was generally more densely expressed in the Abstinence group, compared to both the No Abstinence and Exercise groups. More specifically, out of 33 regions assessed, c-Fos density was significantly greater in the Abstinence group across 18 regions compared to No Abstinence, and 25 regions compared to Exercise. Additionally, a trend towards significance (*p* < 0.100) in the same direction was also found in 8 regions in the No Abstinence group and 3 in the Exercise group. Figure 2 shows c-Fos density in a select set of regions with this typical pattern. Full results can be found in Supplementary Figure S3 & Table S2.

**Figure 2:**
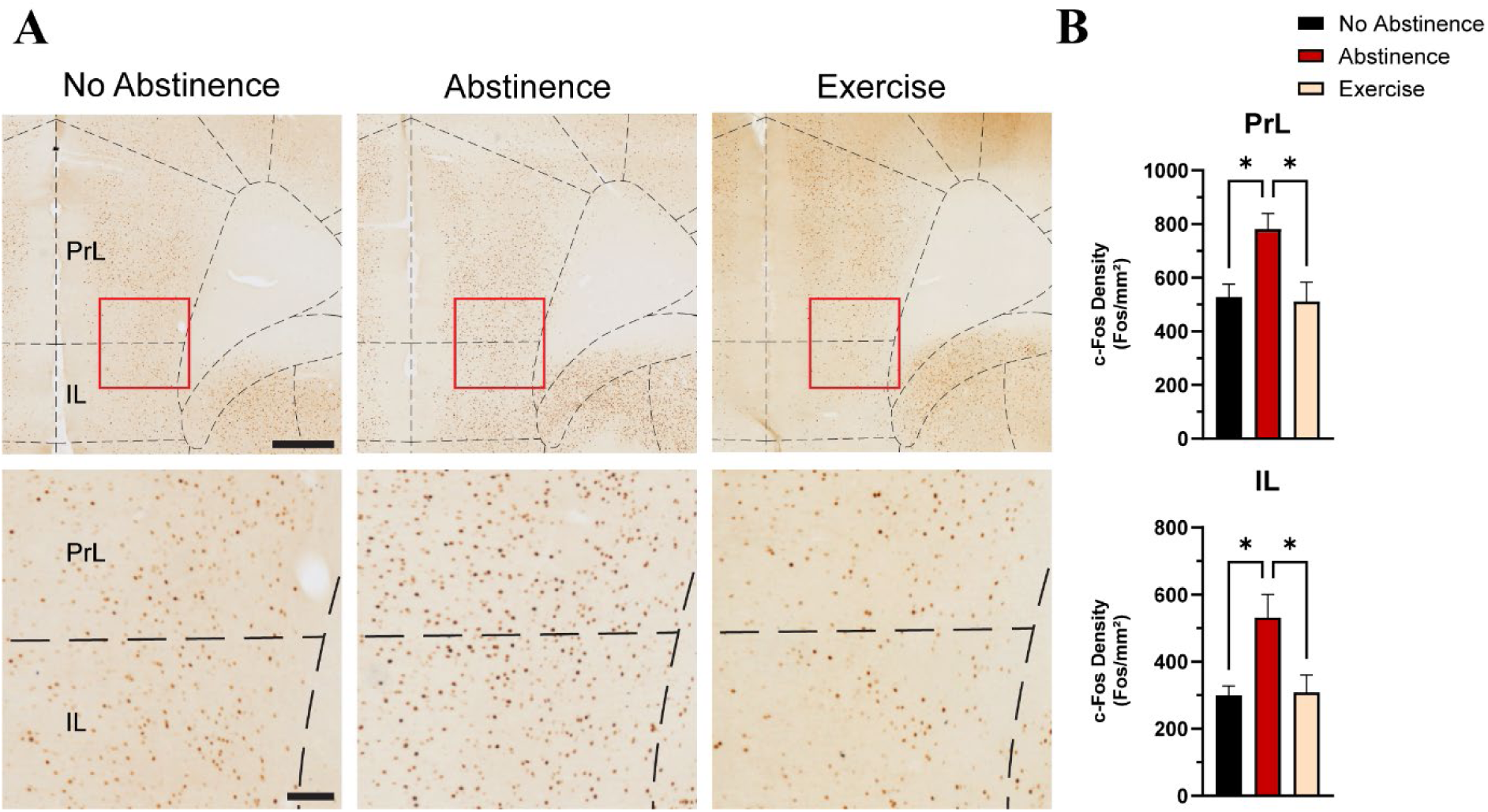
Representative c-Fos Expression. (A) Representative immunohistochemical staining of between-group c-Fos expression in the prelimbic (PrL) and infralimbic (IL) cortex. Scale bars represent 500um (top) and 100um (bottom). Overlay of rat brain atlas plates adapted from Paxinos and Watson (2007). (B) Between-group pattern of c-Fos density in PrL and IL characteristic of the pattern seen across all reward-related regions analysed (Supplementary Figure S3 & Table S2). All data are represented as mean (± SEM). * *p* < 0.05.

##### Hierarchical Clustering

Hierarchical clustering analysis was carried out to identify modules of brain regions that show patterns of highly coordinated activity in each of our abstinence conditions. Each module (Figure 3A – C; red squares) consisted of brain regions that expressed similar overall patterns of Euclidean distances of c-Fos density with all other brain regions (lower Euclidean distance reflected by lighter blue colours). The analysis revealed that the No Abstinence group had a greater number of modules (seven modules) than both the Abstinence (four modules) and Exercise (six modules) groups. Furthermore, this finding was consistent independent of the trim height of the dendrogram (Figure 3D). Of note however, while No Abstinence had more modules than Exercise when trimmed at 50% of the dendrogram height, the number of modules of these two groups was very similar across the whole range of dendrogram height cutoffs. This demonstrates that these two groups retained similar overall modularity profiles.

**Figure 3:**
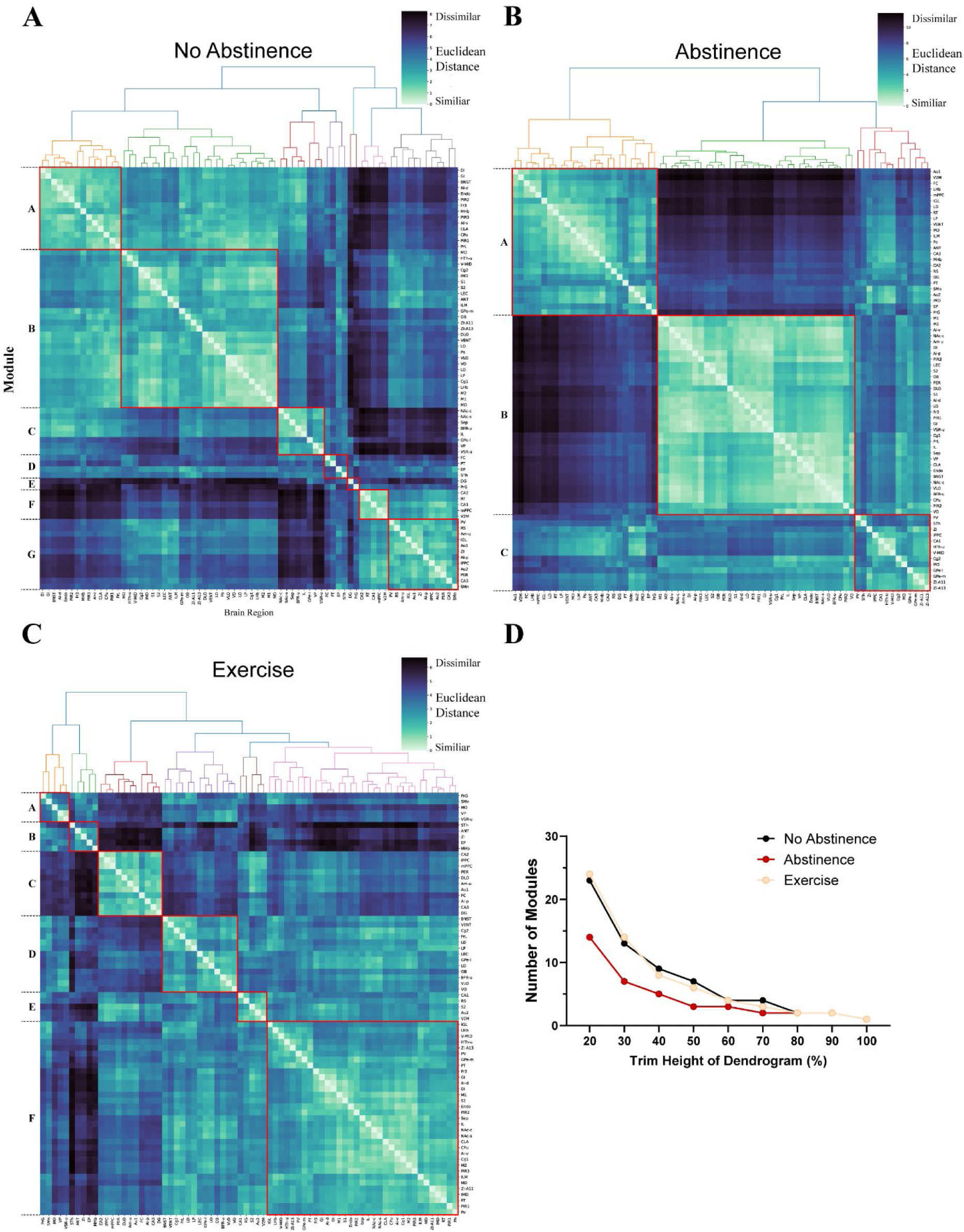
Hierarchical Clustering. (A – C) Hierarchical clustering of Euclidean distance matrix for c-Fos density (c-Fos/mm^2^) of each analysed brain region for the No Abstinence, Abstinence, and Exercise groups. Lighter colours represent smaller Euclidean distances (more similar), darker colours represent greater Euclidean distances (more dissimilar). Red squares indicate modules of brain regions with similar coordinated activity patterns, suggesting functional connectivity. See Table S3 for a full list of brain regions and their modules. (D) Number of modules of coordinated activity across different trim heights of the dendrograms for each group.

In the No Abstinence group, the select reward-related ROIs were distributed across all 7 modules. Module A contained 8 ROIs with similar activation patterns, module B had 10 ROIs, module C had 8, module E had 1, module F had 2, and module G had 4. Interestingly, following Abstinence, a large proportion (21/33) of these regions were grouped in module B, including all of the subregions of the insula and striatum, and most of the orbitofrontal cortex (excluding MO) and the medial prefrontal cortex (excluding Cg2), as well as the amygdala (Am-u), basal forebrain (BFR-u), BNST, Sep, and VP. Of the remaining regions, module A contained the habenula subregions, and the hippocampal subregions CA2, CA3, and DG, while module C contained the subregions of the globus pallidus, as well as Cg2, CA1, medial hypothalamus (HTh-u), MO and PV. Following Exercise, these ROIs were more distributed across modules than after Abstinence, however there was still a larger clustering of certain ROIs in select modules. Most prominently, module F contained 14/33 ROIs, including most of the striatum (excluding the VSR-u) and the insula (excluding the AI-p), half of the medial prefrontal cortex subregions (IL and Cg1) as well as LHb, Sep, GPe-m, HTh-u and PV. Module D had 8 ROIs comprised of most of the orbitofrontal cortex (LO, VO, VLO), the other half of medial prefrontal cortex regions (PrL and Cg2), as well as the BFR-u, BNST, and GPe-l. The remaining ROIs were distributed across the other modules: with hippocampal subregions CA2, CA3, DG, as well as AI-p, Am-u and the DLO in module C; the VP, VSR-u and MO in module A, and only the MHb in module B, and the CA1 in module E. See Table S3 for a full list of brain regions and their modules.

##### Graph Theory

Graph theory analyses were used to further characterise the nature of changes in functional connectivity identified in the hierarchical clustering analysis. Here we focus on changes from No Abstinence to Abstinence and Exercise to examine the connectivity patterns characteristic of each behavioural condition. In No Abstinence (Figure 4A), all modules demonstrated strong average PCs (0.60 – 0.76), suggesting relatively high levels of connectivity across all modules (Table 1). Following Abstinence (Figure 4B), strong average PCs were found for module A and module C, while module B was relatively less interconnected. For the Exercise group (Figure 4C), modules A – E all had strong average PCs, while module F had much weaker inter-module connectivity (see Supplementary Table S3 for a full list of values). This suggests the brain regions in module B of Abstinence, and module F of Exercise, are working more independently than with brain regions contained in other modules.

**Figure 4:**
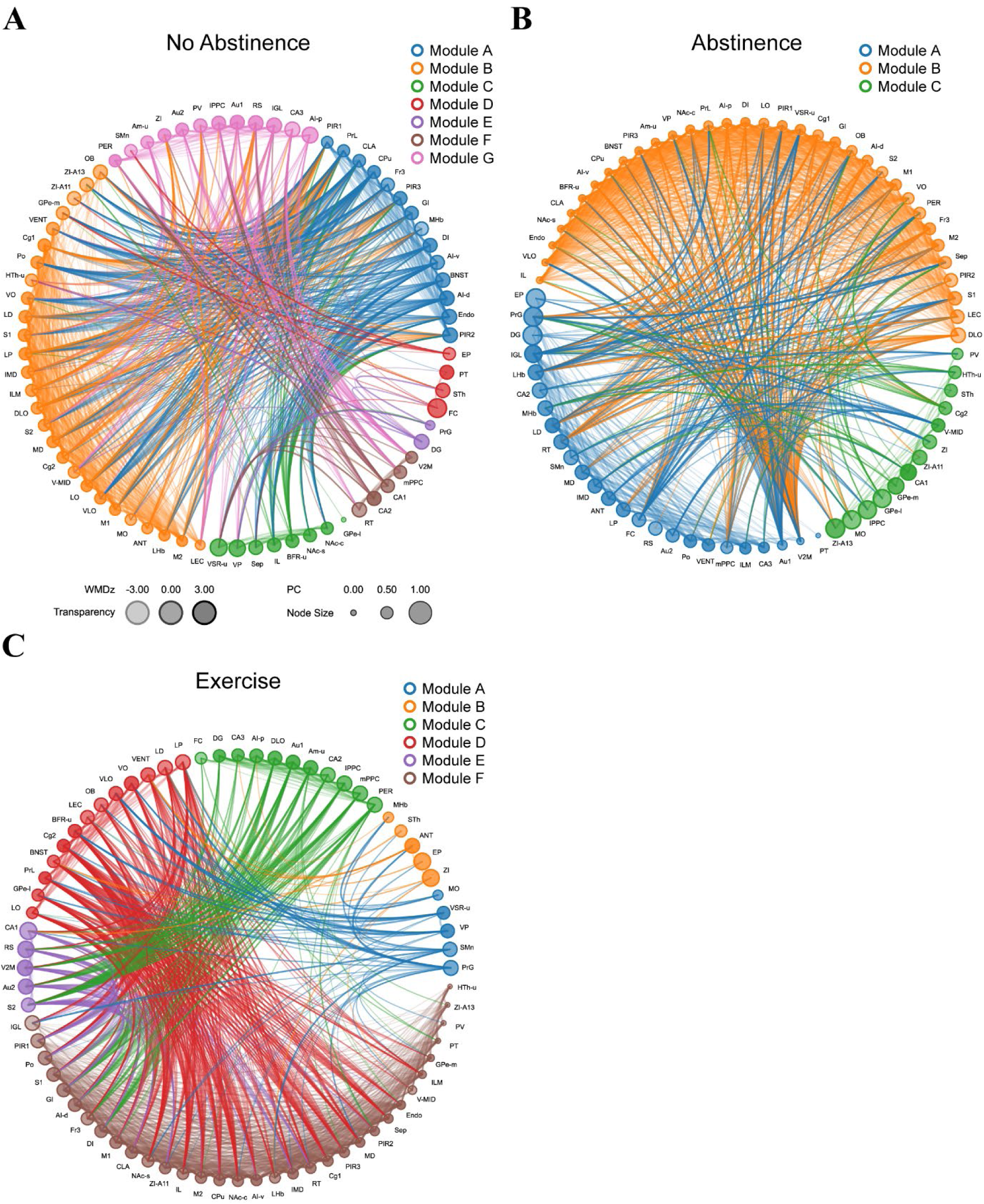
Graph Theory. (A – C) Graphs depict network connectivity for each group. Each node represents a brain region; colour represents hierarchical clustering module, node size represents PC and transparency represents WMDz. Thicker edges represent stronger correlations.

**Table 1.**
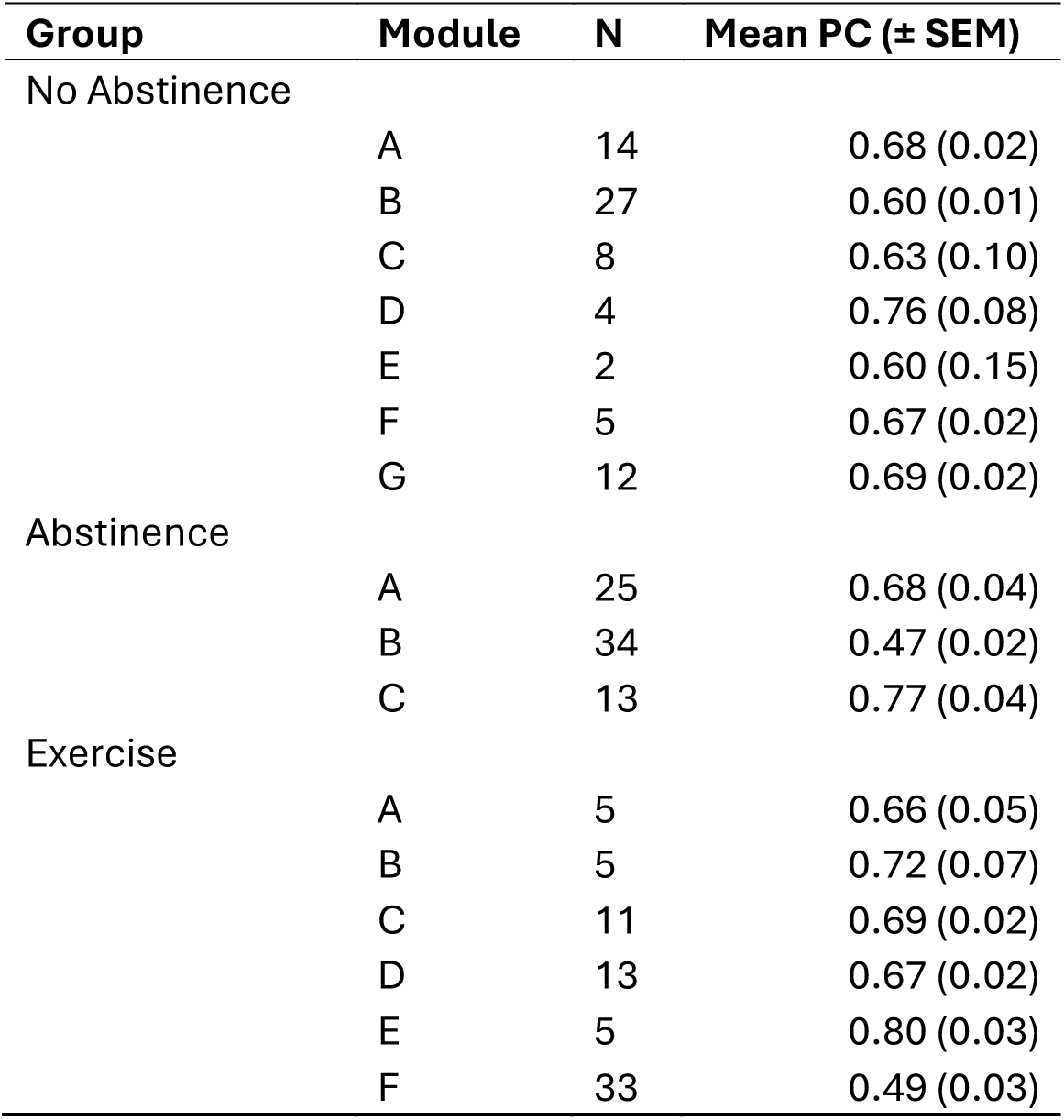
Given the grouping of a large proportion of the select ROIs canonically implicated in reward seeking behaviour in Abstinence module B, we examined which of these regions were acting as provincial hubs (regions with weak PC values and strong WMDz scores) to identify regions that may be driving the independent activity of this module. In Abstinence, these included the AI-v, basal forebrain (BFR-u), caudate putamen (CPu), DI, Endo, GI, NAc-c, PIR3, VP and ventral striatal region (VSR-u). Following Exercise, most of these regions remained clustered together in the less interconnected module F, except for the basal forebrain (BFR-u; in module D), VP and ventral striatal region (VSR-u; in module A). These three regions, that no longer clustered together, interestingly also demonstrated strong WMDz and PC values in their respective clusters after exercise (becoming connector hubs). Hence, while driving coordinated activity across reward-related regions in abstinence, with exercise, the neural activity of these regions becomes more dissimilar to reward-related regions, while acting as connector hubs between modules.

| **Group** | **Module** | **N** | **Mean PC (± SEM)** |
| --- | --- | --- | --- |
| No Abstinence | A | 14 | 0.68 (0.02) |
|  | B | 27 | 0.60 (0.01) |
|  | C | 8 | 0.63 (0.10) |
|  | D | 4 | 0.76 (0.08) |
|  | E | 2 | 0.60 (0.15) |
|  | F | 5 | 0.67 (0.02) |
|  | G | 12 | 0.69 (0.02) |
| Abstinence | A | 25 | 0.68 (0.04) |
|  | B | 34 | 0.47 (0.02) |
|  | C | 13 | 0.77 (0.04) |
| Exercise | A | 5 | 0.66 (0.05) |
|  | B | 5 | 0.72 (0.07) |
|  | C | 11 | 0.69 (0.02) |
|  | D | 13 | 0.67 (0.02) |
|  | E | 5 | 0.80 (0.03) |
|  | F | 33 | 0.49 (0.03) |

## Discussion

The aim of this study was to investigate whether cue-induced reinstatement of alcohol-seeking increased after abstinence (i.e. incubation of craving), whether this effect would be attenuated by exercise, and to assess the associated changes in neural activation and inter-regional coordination with this behaviour. We found that reinstatement was indeed greater when tested 29 days vs 1 day after extinction, and that this increase in reinstatement responding was accompanied by similar increased neural activation in reward circuitry. Voluntary exercise throughout the abstinence period attenuated both reinstatement responding and neural activation in these regions. Interestingly, despite this reduction in neural activation and similar overall modularity profile, Exercise was accompanied by a unique pattern of widespread neural coordination rather than resembling No Abstinence. Together, these results demonstrate that incubation of craving for alcohol-associated cues occurs across a 28-day abstinence period, that exercise attenuates this effect, and for the first time, characterised underlying neural changes involved in the incubation of alcohol craving.

### Abstinence Increases Cue-Induced Alcohol Responding

Our findings are consistent with previous preclinical research investigating the incubation of craving for alcohol (Bienkowski et al., 2004) as well as for other drugs of abuse (Adhikary et al., 2017; Altshuler et al., 2021; Funk et al., 2016; Grimm et al., 2001; Lu et al., 2004; Ma et al., 2014; Markou et al., 2018; Patel & Loweth, 2024; Shalev et al., 2001; Shepard et al., 2004), and reflects changes in craving seen in alcohol-dependent human populations (Bach et al., 2020; Li et al., 2015; Treloar Padovano & Miranda, 2021). These findings support the use of animal models to recapitulate this behavioural phenomenon in the context of alcohol use.

Compared to other studies, there are some differences with respect to the length of abstinence period. We saw evidence of an increase on day 29, while previous reports found incubation on day 56, but not day 28 ((Bienkowski et al., 2004)). The discrepancy may in part reflect the within-session extinction/reinstatement procedure used in that particular study. Rats were tested first under extinction conditions, and the cue was re-introduced in the latter part of the session. At day 28, there was a significant increase in responding during that first “extinction” period, and this non-reinforced responding would likely have influenced subsequent responding when the cue was reintroduced mid-session. On the other hand, at day 56, responding during the initial extinction part of the session was low, leaving the response-outcome association intact and more readily retrieved when the cue was introduced again. It’s also worth noting that in the present study, 7 days of extinction training were conducted prior to the abstinence period, hence rats had in effect undergone 35 days of enforced abstinence (including extinction) prior to test. Therefore, the present finding may reflect the time-dependent increase in craving over the extra week of abstinence. This approach (extinction then abstinence) was adopted to model the time-dependent changes in relapse propensity that may occur throughout abstinence following termination of a structured rehabilitation program where alcohol-seeking is actively inhibited (Perry et al., 2014). Using similar approach, Jupp et al. (2011) found no incubation effect following 5-months of abstinence after 31 days of extinction (5 days a week). Together, these findings suggest that responding for alcohol-associated cues follows an inverted-U shape throughout abstinence, and the incubation of alcohol craving is present 35 days after self-administration.

Regarding alcohol seeking behaviour, it appears that the increase in responding after abstinence is specific to the presentation of a discrete cue. Others have shown that, while maintained across up to 32 days of abstinence, context-induced reinstatement of alcohol-seeking following traditional extinction (Zironi et al., 2006), or punishment-imposed abstinence (Campbell et al., 2019) was not greater at delayed compared to immediate testing. This finding appears to extend across drug classes as cue-induced but not context-induced methamphetamine-seeking increases throughout abstinence (Adhikary et al., 2017). Furthermore, the incubation of craving for cocaine-associated cues is attenuated if the cue is specifically extinguished during the abstinence period (Madsen et al., 2017). This speaks to the importance of the discrete drug-associated cue in the incubation effect. That said, we did not examine time-dependent increases in responding under extinction conditions, and increases in a non-extinguished alcohol-seeking response have been shown after 28 days (Bienkowski et al., 2004). Therefore, it is possible that the observed increase in responding following abstinence may reflect spontaneous recovery. Determining whether such increases in responding are specific to the alcohol-associated cue remains to be elucidated and is an important future direction. Nevertheless, we have demonstrated a clinically relevant finding, whereby increased propensity to seek alcohol at day 29 compared to day 1 of abstinence, despite extinction, may contribute to relapse vulnerability.

### Exercise Attenuates the Incubation of Alcohol Craving

We found that voluntary exercise throughout abstinence prevents time-dependent increases in responding to an alcohol-associated cue. Although we are the first to show this in animals trained to lever press for alcohol, our findings are consistent others who have shown that wheel-running reduces incubation of craving for cocaine (Carroll et al., 2022; Zlebnik & Carroll, 2015). Furthermore, wheel-running also attenuates alcohol and drug-seeking behaviour during other key phases of addiction (for a review, see Carroll (2021)).

A few potential explanations may underlie the protective effect of exercise. One possibility is that the affective disturbances induced by abstinence are reversed or offset by wheel-running access. Indeed, incubation of methamphetamine craving is associated with enhanced trait anxiety and reduced social interaction compared to meth-naïve controls (Everett et al., 2020). Furthermore, abstinence from alcohol consumption in mouse drinking models have demonstrated increased depressive behaviour following 2-weeks of abstinence, which is corrected by wheel-running access throughout this period (Pang et al., 2013). Potentially, such affective disturbances occur in our Abstinence group, but exercise offsets these effects. In addition, social isolation has been shown to impair extinction recall, and exercise prevents such losses (Drummond et al., 2021). Given that extinction occurred prior to the abstinence period in the present study, and animals were socially isolated for 4 hours per day across abstinence, it is possible that exercise may counteract the effects of social isolation on extinction recall. Testing the effect of social isolation and probing potential affective disturbances on relapse propensity remains an important future direction. Another possibility is that exercise may act as an alternative and competing non-drug reward. While in the present procedure there was no choice between exercise or alcohol administration, it has been demonstrated that the wheel running can act as a form of hedonic substitution for alcohol (Booher et al., 2019; Brager & Hammer, 2012; Darlington et al., 2014; Darlington et al., 2016; Ehringer et al., 2009; Gallego et al., 2015; Pichard et al., 2009), and that the reinforcing value of cocaine and wheel-running compete when both are available (Zlebnik et al., 2010). Hence, exercise interventions may provide a rewarding alternative that reduces the incentive salience of drug-associated cues throughout abstinence when the opportunity to consume alcohol arises.

The protective effect of exercise also parallels findings regarding environmental enrichment, an intervention to improve animal wellbeing by increasing the complexity of their living environments (Solinas et al., 2021). Enriched environments reduce incubation of craving across drug classes (Chauvet et al., 2012; Sikora et al., 2018; Thiel et al., 2012; Thiel et al., 2010) and for non-drug rewards (Grimm et al., 2016; Grimm et al., 2025). The effect of wheel-running could be considered a form of environmental enrichment, working similarly to reduce craving. Together, this supports a potential role for adjunctive lifestyle interventions throughout abstinence to reduce relapse propensity.

### Neural Activation Associated with The Incubation of Alcohol Craving

We used c-Fos immunoreactivity to assess the neural correlates associated with incubation of alcohol craving. Initial analysis showed that the between-group pattern of activity across the reward circuitry mirrored the pattern observed in behaviour – i.e. for almost all regions assessed, c-Fos immunoreactivity was significantly greater in the abstinence condition compared to both no abstinence and exercise conditions. These findings are consistent with related studies demonstrating increased neural recruitment in reward-related regions for the incubation of craving for other drug and non-drug rewards (Altshuler et al., 2021; Caprioli et al., 2017; Davis et al., 2021; Fanous et al., 2012; Funk et al., 2016; Grimm et al., 2016; Li et al., 2018; Rossi et al., 2020).

Only one other study has provided a comprehensive assessment of neural correlates in a similar preparation to ours – that is, cue induced reinstatement of alcohol-seeking after extinction or extinction plus abstinence (Jupp et al. 2011). In this study, no behavioural evidence of incubation of craving was found. Interestingly, however, despite responding being equivalent, rats tested after abstinence did show greater c-Fos expression in the medial prefrontal, piriform and orbitofrontal cortices compared to those tested immediately after extinction. As discussed previously, the longer 5-month abstinence period in this study may capture a later point on an inverted U-shaped curve in cue-reactivity. This suggests that there are ongoing abstinence-associated changes in neural recruitment, even if this does not manifest in increased alcohol-seeking. In contrast, in the present study, following 28 days of abstinence we similarly observed increased activity across the frontal cortex, but also in the striatum, coincident with an increase in cue-elicited alcohol-seeking. Indeed, strengthening of corticostriatal circuits underpin incubation of psychostimulant and opioid craving (Altshuler et al., 2021; Ma et al., 2014), and together these findings suggest that with ongoing abstinence there may be recovery of neural changes that manifests in decreased co-activation of corticostriatal circuits and consequent reduction of relapse propensity.

Although this hypothesis could not be tested in the current study, we were able to use hierarchical clustering and graph theory analyses to explore functional connectivity between brain regions pre and post abstinence. In the No Abstinence group, seven different modules of brain regions were identified as clusters with coordinated neural activity. Interestingly, many of the reward-seeking related ROIs were spread across these modules, indicating a relatively uncoordinated neural response to alcohol-associated cues. Similarly, graph theory analyses indicated strong intermodular connectivity, as indexed by all modules displaying high levels of uniformity in their connections with each other (all PCs > 0.5). After a period of abstinence, a vast majority of the select ROIs were grouped in the same module (module B) accompanied by uniquely weak intermodular connectivity (PC < 0.5). In other words, neural activity of the brain regions in this cluster were more coordinated with each other, and less coordinated with regions in other clusters.

The coordination of cortical, amygdala and striatal regions in this module is consistent with the strengthening of corticostriatal and amygdala-striatal circuits throughout abstinence observed with incubation of psychostimulant rewards (Altshuler et al., 2020; Lee et al., 2013; Ma et al., 2014; Wolf, 2016, 2025). Such enhanced, independent coordination of these regions may underscore the increase in relapse observed – implying a ‘Reinstatement Cluster’ of brain regions after abstinence. Furthermore, this cluster included the insula, orbitofrontal, and medial prefrontal cortex, which are key regions associated with the “preoccupation/anticipation” (craving) stage of addiction (Koob & Volkow, 2016). While these findings represent an early exploratory approach to understanding the neural mechanisms associated with the incubation of craving for alcohol associated cues, they point to a common mechanism underlying increased craving and subsequent relapse across drug classes.

Following exercise, we found that overall neural activation returned to a similar modularity profile of clustered regions as the no abstinence condition. Nevertheless, many of the regions in the Reinstatement Cluster remained clustered in module F (see Fig 4 and supplementary table S3), which also displayed a similar modular independence (high intra- and low inter- modular connectivity). However, it is worth noting that certain key ROIs were no longer coordinated after exercise treatment. These brain regions represent candidate neural loci where exercise may be exerting effects to reduce relapse propensity after abstinence. For example, the posterior insula (AI-p), amygdala (Am-u), basal forebrain (BFR-u), BNST, dorsolateral-, lateral-, ventrolateral-, and lateral-orbitofrontal cortex (DLO, LO, VLO, VO), prelimbic cortex (PrL), ventral pallidum (VP), and ventral striatum (VSR-u) all shifted to clusters with stronger inter-modular connectivity – indicating reduced rigidity in their coordination across the all regions analysed. Graph theory analyses identified the VP and VSR-u as potential regions driving the coordination of the Reinstatement Cluster after abstinence; subsequently they shifted into their own module with exercise treatment, while retaining strong interconnectedness with other reinstatement related regions. Given the limitations of the current version of the WHS atlas, it is difficult to make conclusions about the broader VSR-u. However, the ‘VSR-u’ delineation overlaps with the lateral nucleus accumbens shell and the IPAC, which play a role in reward consumption, motivated behaviour, and integration of reward and context memory (Chen et al., 2023; Liu et al., 2023; Yang et al., 2018). In addition, the VP, while not studied in the context of the incubation of craving, is considered the final output region in the ‘final common pathway’ of relapse to drug seeking (Kalivas & Volkow, 2005; Koob & Volkow, 2010) and is particularly implicated in the reinstatement of alcohol seeking (Perry & McNally, 2013; Prasad & McNally, 2020) and encoding incentive motivation for rewarding stimuli (Smith et al., 2009). Hence, exercise treatment may be working to disengage the VP and parts of ventral striatum from the broader reinstatement circuitry, to prevent increased relapse behaviour.

Alternatively, the lack of increased relapse after exercise may result from the disentanglement of coordinated activity between the subregions of the medial prefrontal and orbitofrontal cortices, that underpin the ‘craving’ stage of addiction (Koob & Volkow, 2016), as well as the striatum and amygdala – as these connections are known to drive the incubation of psychostimulant and opioid craving (Altshuler et al., 2020; Lee et al., 2013; Ma et al., 2014; Wolf, 2016, 2025). Furthermore, the exercise-induced downstream “decoupling” of corticostriatal and amygdala-striatal coordination resembles the hypothesis described earlier regarding the differences seen in brain activity patterns across longer periods of alcohol abstinence. While no specific neural mechanisms can be solely determined from the present approach, these findings provide evidence that exercise may be acting upon select brain regions or pathways to exert relapse protective effects by restoring widespread interconnectivity.

In summary, we showed that abstinence increases the incentive motivation of alcohol-associated cues, consistent with incubation of craving. This effect was significantly attenuated if rats were allowed to exercise across abstinence. Increased alcohol-seeking was associated with increased brain activity in reward circuits, decreased modularity of neural activation, and enhanced coordination of reward-related circuitry. Importantly, exercise treatment throughout this period prevented this change in modularity and resulted in specific changes in coordination of brain regions implicated in the expression of relapse behaviour and the incubation of craving for psychostimulant and opioid reward. These regions represent candidate neural loci whereby exercise may be exerting neural adaptations to prevent the increase in relapse propensity. This is the first study to characterise neural activation associated with the incubation of craving for alcohol-associated cues and provides a neural basis for exercise treatment to help prevent relapse to alcohol seeking.

## Acknowledgements

We thank the Central Animal Facility staff at Macquarie University for animal husbandry.

## Contributions

TMF and CJP designed the experiment and prepared the manuscript. TMF performed all experiment, with support from AOC, MS, AIJK & LU. AIJK provided particular support for graph theory analysis. CJP acquired funds for the research. All authors reviewed and edited the manuscript.

## Data availability statement

The data that support the findings of this study are available from the corresponding author upon reasonable request.

