## Supplementary materials for "Incubation of craving for alcohol-associated cues is reduced by running-wheel exercise"

### Materials and Methods

#### *QUINT Workflow*

Images of coronal sections from Bregma +4.68mm To Bregma -4.20mm were processed using the QUINT workflow. Prior to quantification, images were pre-processed to be downscaled to a computationally feasible resolution, crop out folded areas of tissue, rotated to be upright, and to determine the serial order from anterior to posterior. Furthermore, specific images with extreme tissue distortion or damage were excluded.

Sections were anchored in 3D space to the Waxholm Space atlas of the rat brain (WHS atlas) version 4 (Kleven et al., 2023), using QuickNII, which accounts for the angle of sectioning (Figure S1A). VisuAlign was utilised to make further non-linear adjustments to the atlas overlay to fit the histological sections more accurately, and to account for tissue distortion due to immunohistochemistry and slide-mounting procedures (Figure S1B).

To identify c-Fos expression from background tissue, a pixel classification workflow in Ilastik was employed. The machine learning model (a Random Forest classifier) was trained using a representative sample of 48 sections: 4 sections spanning the anterior/posterior axis, from 4 brains for each No Abstinence, Abstinence and Exercise. Labels were assigned to 'c-Fos' or 'background' to annotate pixels on the images that represented c-Fos staining, or did not (e.g. background tissue, or where no tissue was present; Figure S1C), respectively. All pixel feature types (colour/intensity, edge, and texture) were selected to discern label classification. User annotations were iteratively made across all 48 sections, ensuring inclusion of different staining appearances, until the machine learning predictions were deemed to accurately identify c-Fos across all the tissue used for training. The classifier predictions were then applied to all sections of each brain via batch processing to produce a segmentation image for each section. A watershed function was then applied to each segmentation image to separate overlapping c-Fos cells in the FIJI software (Schindelin et al., 2012) to produce the final c-Fos segmentation images for quantification.

The Nutil software 'Quantifier' operation was used to combine the VisuAlign atlas maps with the c-Fos segmentation images to quantify c-Fos expression in each brain region.

***Figure S1: QUINT Workflow Overview***

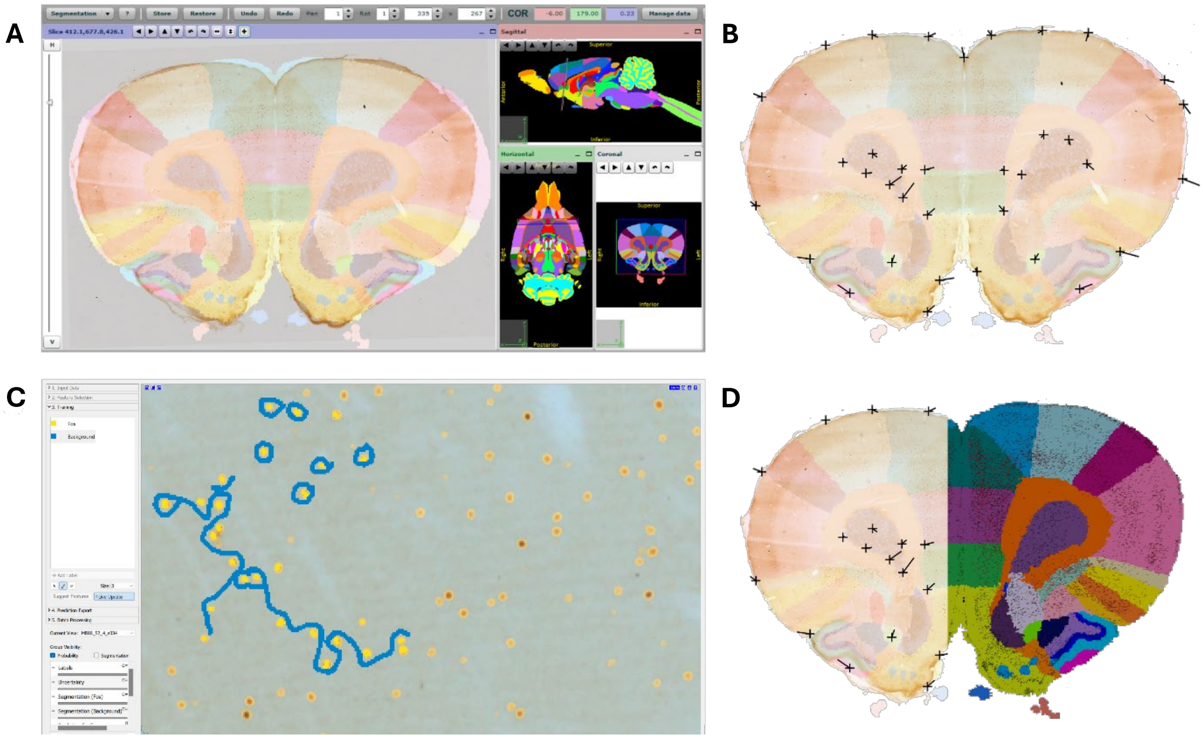


An overview of the software used in the QUINT workflow, using a representative section. (A) QuickNII software allows for histological sections to be anchored in 3D space using the Waxholm Space atlas of the rat brain version 4, accounting for the angle of sectioning. (B) VisuAlign allows for adjustment of the atlas overlay to account for tissue distortion throughout histological procedures. Black crosses represent where the atlas overlay has been moved, to align with the histological section. (C) Ilastik is used to train a machine learning model to classify whether each pixel on the photomicrograph represents c-Fos (yellow annotations), or background (blue annotations). Transparent yellow and blue overlay represents the model's predictions based on all user-defined annotations to all non-defined pixels. (D) Figure depicts the how the final atlas overlay (left) is combined with the Ilastik procedure output, to quantify c-Fos cells identified by the machine learning model in the respective region of the atlas (right).

### *Table S1: Brain Regions of Fos Quantification*

| Region name | Abbreviations | WHS Region ID |
|---|---|---|
| Agranular insular cortex dorsal area | AI-d | 410 |
| Agranular insular cortex posterior area | AI-p | 424 |
| Agranular insular cortex ventral area | AI-v | 409 |
| Amygdaloid area unspecified | Am-u | 501 |
| Basal forebrain region unspecified | BFR-u | 82 |
| Bed nucleus of the stria terminalis | BNST | 93 |
| Caudate putamen | CPu | 197 |
| Cingulate area 1 | Cg1 | 411 |
| Cingulate area 2 | Cg2 | 10 |
| Claustrum | CLA | 412 |
| Cornu ammonis 1 | CA1 | 98 |
| Cornu ammonis 2 | CA2 | 97 |
| Cornu ammonis 3 | CA3 | 95 |
| Dentate gyrus | DG | 96 |
| Dorsolateral orbital area | DLO | 404 |
| Dysgranular insular cortex | DI | 414 |
| Endopiriform nucleus | Endo | 500 |
| Entopeduncular nucleus | EP | 32 |
| Fasciola cinereum | FC | 99 |
| Frontal association area 3 | Fr3 | 407 |
| Globus pallidus external lateral part | GPe-l | 198 |
| Globus pallidus external medial part | GPe-m | 195 |
| Granular insular cortex | GI | 416 |
| Hypothalamic region unspecified | HTh-u | 48 |
| Infralimbic area | IL | 413 |
| Intergeniculate leaflet | IGL | 272 |
| Intermediodorsal thalamic nucleus | IMD | 260 |
| Lateral entorhinal cortex | LEC | 115 |
| Lateral habenular nucleus | LHb | 206 |
| Lateral orbital area | LO | 401 |
| Medial habenular nucleus | MHb | 207 |
| Medial orbital area | MO | 403 |
| Nucleus accumbens core | NAc-c | 184 |
| Nucleus accumbens shell | NAc-s | 192 |
| Nucleus of the stria medullaris | SMn | 81 |
| Parataenial thalamic nucleus | PT | 211 |
| Paraventricular thalamic nuclei (anterior and posterior) | PV | 242 |
| Parietal association cortex lateral area | IPPC | 432 |
| Parietal association cortex medial area | mPPC | 433 |
| Piriform cortex layer 1 | PIR1 | 181 |
| Piriform cortex layer 2 | PIR2 | 182 |
| Piriform cortex layer 3 | PIR3 | 183 |
| Posterior thalamic nucleus | Po | 230 |
| Pregeniculate nucleus | PrG | 204 |
| Prelimbic area | PrL | 405 |

| | | | |
|---|---|---|---|
| Primary auditory area | | Au1 | 151 |
| Primary motor area | | M1 | 408 |
| Secondary motor area | | M2 | 406 |
| Secondary somatosensory area | | S2 | 422 |
| Secondary visual area medial part | | V2M | 448 |
| Septal region | | Sep | 40 |
| Subthalamic nucleus | | STh | 3 |
| Ventral orbital area | | VO | 402 |
| Ventral pallidum | | VP | 193 |
| Ventral striatal region unspecified | | VSR-u | 199 |
| Ventrolateral orbital area | | VLO | 400 |
| Zona incerta A11 dopamine cells | | ZI-A11 | 284 |
| Zona incerta A13 dopamine cells | | ZI-A13 | 238 |

**Grouped Regions**

| **Grouped Region Name** | **Subregion** | | |
|---|---|---|---|
| Olfactory bulb | | OB | 1020 |
| | Nucleus of the lateral olfactory tract | | 502 |
| | Olfactory bulb unspecified | | 66 |
| Perirhinal cortex | | PER | 1024 |
| | Perirhinal area 35 | | 112 |
| | Perirhinal area 36 | | 113 |
| Reticular (pre)thalamic nucleus | | RT | 1033 |
| | Reticular (pre)thalamic nucleus auditory segment | | 164 |
| | Reticular (pre)thalamic nucleus unspecified | | 200 |
| Retrosplenial cortex | | RS | 1058 |
| | Retrosplenial dysgranular area | | 427 |
| | Retrosplenial granular area | | 430 |
| Primary somatosensory area | | S1 | 1067 |
| | Primary somatosensory area barrel field | | 425 |
| | Primary somatosensory area dysgranular zone | | 418 |
| | Primary somatosensory area face representation | | 420 |
| | Primary somatosensory area forelimb representation | | 417 |
| | Primary somatosensory area hindlimb representation | | 423 |
| | Primary somatosensory area trunk representation | | 429 |
| Secondary auditory area | | Au2 | 1073 |
| | Secondary auditory area dorsal part | | 152 |
| | Secondary auditory area ventral part | | 153 |
| Anterior nuclei of the dorsal thalamus | | ANT | 1085 |
| | Anterodorsal thalamic nucleus | | 213 |
| | Anteromedial thalamic nucleus | | 254 |
| | Anteroventral thalamic nucleus dorsomedial part | | 214 |
| | Anteroventral thalamic nucleus ventrolateral part | | 215 |
| | Interanteromedial thalamic nucleus | | 255 |
| Ventral midline group of the dorsal thalamus | | V-MID | 1088 |
| | Retroreuniens thalamic nucleus | | 268 |
| | Reuniens thalamic nucleus | | 219 |
| | Rhomboid thalamic nucleus | | 216 |

| | | | |
|---|---|---|---|
| | Xiphoid thalamic nucleus | | 218 |
| Mediodorsal nucleus of the dorsal thalamus | | MD | 1089 |
| | Mediodorsal thalamic nucleus central part | | 233 |
| | Mediodorsal thalamic nucleus lateral part | | 232 |
| | Mediodorsal thalamic nucleus medial part | | 240 |
| Ventral nuclei of the dorsal thalamus | | VENT | 1090 |
| | Angular thalamic nucleus | | 223 |
| | Submedius thalamic nucleus | | 222 |
| | Ventral anterior thalamic nucleus | | 293 |
| | Ventral posterior nucleus of the thalamus parvicellular part | | 266 |
| | Ventral posterolateral thalamic nucleus | | 294 |
| | Ventral posteromedial thalamic nucleus | | 227 |
| | Ventrolateral thalamic nucleus | | 231 |
| | Ventromedial thalamic nucleus | | 221 |
| Intralaminar nuclei of the dorsal thalamus | | ILM | 1092 |
| | Central lateral thalamic nucleus | | 248 |
| | Central medial thalamic nucleus | | 247 |
| | Paracentral thalamic nucleus | | 246 |
| Lateral posterior (pulvinar) complex of the dorsal thalamus | | LP | 1094 |
| | Lateral posterior thalamic nucleus lateral part | | 283 |
| | Lateral posterior thalamic nucleus mediorostral part | | 285 |
| Laterodorsal thalamic nuclei of the dorsal thalamus | | LD | 1095 |
| | Dorsal lateral geniculate nucleus | | 205 |
| | Laterodorsal thalamic nucleus dorsomedial part | | 228 |
| | Laterodorsal thalamic nucleus ventrolateral part | | 229 |
| Zona incerta | | ZI | 1082 |
| | Zona incerta dorsal part | | 235 |
| | Zona incerta rostral part | | 257 |
| | Zona incerta ventral part | | 236 |

#### *Graph Theory Metrics*

For the WMDz to represent a z-score of intra-module connectivity of node $i$ let:

$$WMDz_i = \frac{k_i - \bar{k}_{s_i}}{\sigma_{k_{s_i}}}$$

Where $k_i$ is the summed weight of all edges with other nodes within its own module $s_i$; $\bar{k}_{s_i}$ is the average of the summed weight of all nodes $k$ edges within in the same module ($s_i$), and $\sigma_{k_{s_i}}$ is the standard deviation of $k$ in $s_i$. Hence, the WMDz represents the normalised strength of a given brain region's connection to its own module (higher WMDz), or other modules (lower WMDz). Values > 0.75 were considered strong WMDz scores.

To determine the uniformity of the distribution of edges amongst all modules, let the PC of node $i$ be:

$$P_i = 1 - \sum_{s=1}^{N_M} \left(\frac{k_{is}}{k_i}\right)^2$$

Where $k_{is}$ represents the summed weight of all edges between node $i$ and all nodes in module $s$, and $k_i$ is the summed weight of all edges between node $i$ and all other nodes in the network. As the highest possible PC is determined by the number of modules (e.g. max value of 2/3 for 3 modules), PC values were normalised (to range from 0 to 1) by dividing the PC value by the theoretical maximum value, to allow for comparison between groups. Hence, the PC represents a measure of inter-module connectivity, where a value of 0 represents that the brain region has connections only to other regions in its own module, and value of 1 if its connections are uniformly distributed across all modules. Values greater than 0.5 were considered strong PC scores.

**Results**

***Figure S2: Distance Run (m) in Exercise***

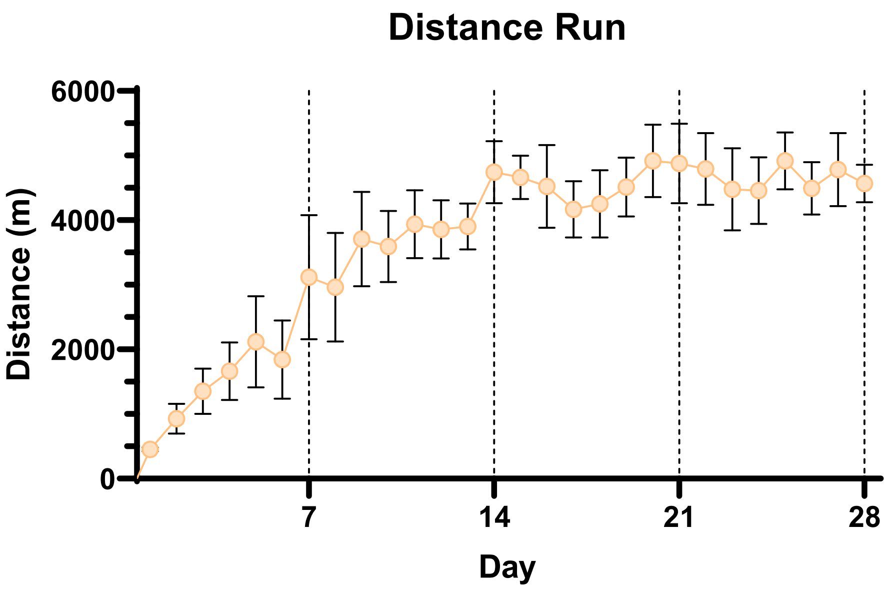


Mean distance (m) ± SEM run by the Exercise group throughout abstinence

### *Figure S3: c-Fos Density of Select ROIs*

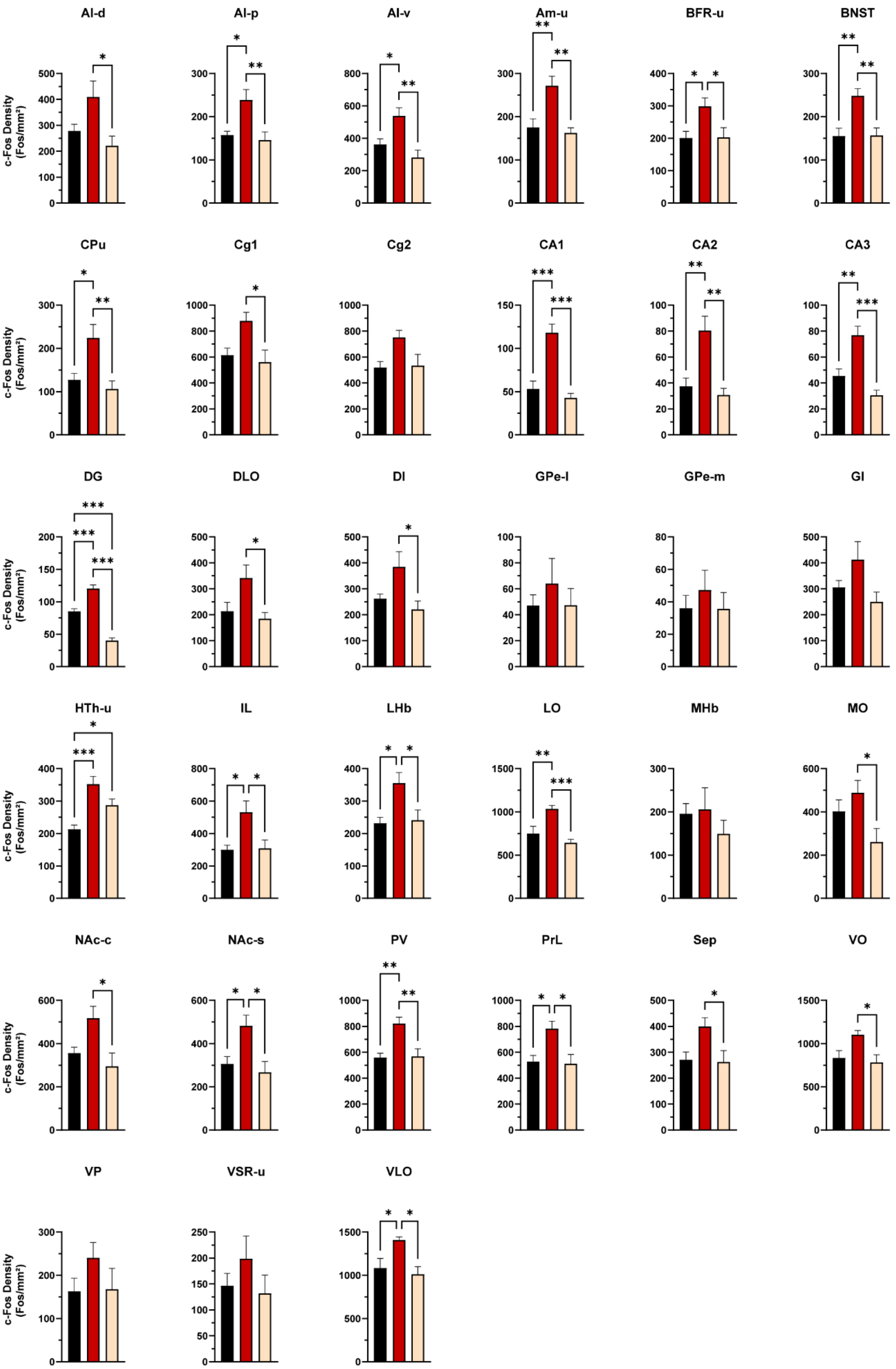


### *Table S2: ANOVAs of Select ROIs*

| ROI | *F*-value | *p*-value | $R^2$ | Between Group Comparisons (*p*-value) | | |
|---|---|---|---|---|---|---|
| | | | | No Abstinence vs. Abstinence | No Abstinence vs. Exercise | Abstinence vs. Exercise |
| AI-d | $F_{2, 15}$ = 4.79 | 0.025 | 0.39 | 0.123 | 0.339 | 0.022 |
| AI-p | $F_{2, 15}$ = 7.43 | 0.006 | 0.50 | 0.019 | 0.902 | 0.008 |
| AI-v | $F_{2, 15}$ = 8.71 | 0.003 | 0.54 | 0.034 | 0.431 | 0.003 |
| Am-u | $F_{2, 15}$ = 10.58 | 0.001 | 0.59 | 0.005 | 0.881 | 0.002 |
| BFR-u | $F_{2, 15}$ = 4.73 | 0.026 | 0.39 | 0.042 | 0.998 | 0.047 |
| BNST | $F_{2, 15}$ = 9.29 | 0.002 | 0.55 | 0.005 | 0.997 | 0.006 |
| CPu | $F_{2, 15}$ = 7.65 | 0.005 | 0.50 | 0.022 | 0.798 | 0.006 |
| Cg1 | $F_{2, 15}$ = 5.50 | 0.016 | 0.42 | 0.052 | 0.864 | 0.019 |
| Cg2 | $F_{2, 15}$ = 3.98 | 0.041 | 0.35 | 0.057 | 0.984 | 0.079 |
| CA1 | $F_{2, 15}$ = 24.09 | <0.001 | 0.76 | <0.001 | 0.658 | <0.001 |
| CA2 | $F_{2, 15}$ = 11.53 | 0.001 | 0.61 | 0.004 | 0.822 | 0.001 |
| CA3 | $F_{2, 15}$ = 17.77 | <0.001 | 0.70 | 0.003 | 0.179 | <0.001 |
| DG | $F_{2, 15}$ = 76.27 | <0.001 | 0.91 | <0.001 | <0.001 | <0.001 |
| DLO | $F_{2, 15}$ = 4.83 | 0.024 | 0.39 | 0.074 | 0.858 | 0.027 |
| DI | $F_{2, 15}$ = 4.52 | 0.029 | 0.38 | 0.110 | 0.753 | 0.029 |
| GPe-l | $F_{2, 15}$ = 0.47 | 0.637 | 0.06 | 0.684 | 0.999 | 0.691 |
| GPe-m | $F_{2, 15}$ = 0.42 | 0.664 | 0.05 | 0.721 | 0.999 | 0.703 |
| GI | $F_{2, 15}$ = 2.96 | 0.083 | 0.28 | 0.285 | 0.700 | 0.073 |
| HTh-u | $F_{2, 15}$ = 13.54 | <0.001 | 0.64 | <0.001 | 0.035 | 0.070 |
| IL | $F_{2, 15}$ = 6.31 | 0.010 | 0.46 | 0.018 | 0.993 | 0.022 |
| LHb | $F_{2, 15}$ = 6.05 | 0.012 | 0.45 | 0.018 | 0.971 | 0.028 |
| LO | $F_{2, 15}$ = 12.15 | <0.001 | 0.62 | 0.009 | 0.417 | 0.001 |
| MHb | $F_{2, 15}$ = 0.67 | 0.527 | 0.08 | 0.980 | 0.655 | 0.538 |
| MO | $F_{2, 15}$ = 3.99 | 0.041 | 0.35 | 0.550 | 0.225 | 0.034 |
| NAc-c | $F_{2, 15}$ = 5.29 | 0.018 | 0.41 | 0.089 | 0.670 | 0.017 |
| NAc-s | $F_{2, 15}$ = 6.46 | 0.010 | 0.46 | 0.037 | 0.815 | 0.011 |
| PV | $F_{2, 15}$ = 9.52 | 0.002 | 0.56 | 0.004 | 0.991 | 0.006 |
| PrL | $F_{2, 15}$ = 6.28 | 0.010 | 0.46 | 0.025 | 0.978 | 0.017 |
| Sep | $F_{2, 15}$ = 4.42 | 0.031 | 0.37 | 0.062 | 0.985 | 0.045 |

| | | | | | | |
|---|---|---|---|---|---|---|
| VO | $F_{2, 15} = 5.04$ | 0.021 | 0.40 | 0.062 | 0.883 | 0.025 |
| VP | $F_{2, 15} = 1.26$ | 0.311 | 0.14 | 0.357 | 0.996 | 0.401 |
| VSR-u | $F_{2, 15} = 1.00$ | 0.391 | 0.12 | 0.556 | 0.954 | 0.393 |
| VLO | $F_{2, 15} = 6.08$ | 0.012 | 0.45 | 0.042 | 0.830 | 0.013 |

Pairwise between-group comparison p-values are Tukey's HSD adjusted

***Table S3: Hierarchical Clustering Modules and Graph Theory Metrics***

| Group | Module | Brain Region | WMDz | PC |
|---|---|---|---|---|
| No Abstinence | | | | |
| | A | AI-d | 1.19 | 0.75 |
| | | DI | 0.73 | 0.70 |
| | | PIR2 | 0.72 | 0.75 |
| | | CPu | 0.66 | 0.65 |
| | | Endo | 0.63 | 0.75 |
| | | PIR3 | 0.53 | 0.68 |
| | | GI | 0.30 | 0.68 |
| | | BNST | 0.19 | 0.74 |
| | | CLA | 0.02 | 0.63 |
| | | PrL | -0.03 | 0.60 |
| | | AI-v | -0.50 | 0.72 |
| | | PIR1 | -0.59 | 0.58 |
| | | Fr3 | -1.12 | 0.67 |
| | | MHb | -2.73 | 0.69 |
| | B | LP | 1.06 | 0.60 |
| | | LO | 0.91 | 0.56 |
| | | Cg1 | 0.86 | 0.63 |
| | | LHb | 0.86 | 0.53 |
| | | MD | 0.85 | 0.59 |
| | | M2 | 0.84 | 0.52 |
| | | Cg2 | 0.79 | 0.59 |
| | | M1 | 0.78 | 0.55 |
| | | V-MID | 0.72 | 0.56 |
| | | S1 | 0.66 | 0.61 |
| | | IMD | 0.65 | 0.60 |
| | | LD | 0.63 | 0.61 |
| | | VO | 0.55 | 0.62 |
| | | Po | 0.53 | 0.62 |
| | | S2 | 0.40 | 0.59 |
| | | ILM | 0.15 | 0.60 |
| | | DLO | 0.14 | 0.59 |
| | | ANT | -0.20 | 0.54 |
| | | VLO | -0.24 | 0.64 |
| | | V2M | -0.49 | 0.58 |
| | | LEC | -0.80 | 0.47 |
| | | GPe-m | -1.25 | 0.66 |

| | | | | |
|---|---|---|---|---|
| | | MO | -1.31 | 0.55 |
| | | HTh-u | -1.44 | 0.62 |
| | | OB | -1.50 | 0.72 |
| | | ZI-A13 | -1.82 | 0.70 |
| | | ZI-A11 | -2.34 | 0.68 |
| | C | BFR-u | 0.60 | 0.66 |
| | | NAc-s | 0.60 | 0.65 |
| | | NAc-c | 0.51 | 0.61 |
| | | IL | 0.46 | 0.69 |
| | | VP | 0.36 | 0.85 |
| | | Sep | -0.01 | 0.71 |
| | | VSR-u | -0.14 | 0.89 |
| | | GPe-l | -2.38 | 0.00 |
| | D | PT | 1.22 | 0.70 |
| | | FC | 0.09 | 0.97 |
| | | STh | -0.09 | 0.73 |
| | | EP | -1.22 | 0.62 |
| | E | DG | NA | 0.75 |
| | | PrG | NA | 0.45 |
| | | CA2 | 1.52 | 0.72 |
| | | VENT | 0.06 | 0.60 |
| | | mPPC | -0.13 | 0.64 |
| | | CA1 | -0.15 | 0.67 |
| | | RT | -1.29 | 0.73 |
| | F | Au1 | 1.27 | 0.71 |
| | | PER | 1.14 | 0.61 |
| | | RS | 0.69 | 0.74 |
| | | ZI | 0.55 | 0.64 |
| | | lPPC | 0.47 | 0.69 |
| | | IGL | 0.42 | 0.76 |
| | | Au2 | 0.36 | 0.64 |
| | | PV | -0.39 | 0.66 |
| | | AI-p | -0.42 | 0.82 |
| | | Am-u | -0.63 | 0.63 |
| | | CA3 | -1.67 | 0.77 |
| | | SMn | -1.81 | 0.63 |
| Abstinence | | | | |
| | A | LD | 1.48 | 0.75 |
| | | IGL | 1.36 | 0.90 |
| | | ILM | 1.19 | 0.57 |
| | | VLO | 0.96 | 0.63 |
| | | Po | 0.85 | 0.64 |
| | | CA3 | 0.85 | 0.52 |
| | | MD | 0.81 | 0.73 |
| | | LHb | 0.81 | 0.78 |
| | | ANT | 0.77 | 0.71 |
| | | IMD | 0.63 | 0.71 |
| | | LP | 0.59 | 0.70 |

| | | | |
|---|---|---|---|
| | RT | 0.55 | 0.74 |
| | MHb | 0.22 | 0.76 |
| | PrG | -0.15 | 0.99 |
| | Au2 | -0.34 | 0.66 |
| | CA2 | -0.46 | 0.77 |
| | Au1 | -0.63 | 0.46 |
| | FC | -0.64 | 0.69 |
| | SMn | -0.70 | 0.74 |
| | mPPC | -0.96 | 0.60 |
| | VENT | -1.02 | 0.29 |
| | DG | -1.38 | 0.99 |
| | EP | -1.55 | 0.99 |
| | RS | -1.62 | 0.68 |
| | PT | -1.62 | 0.00 |
| B | BFR-u | 1.22 | 0.32 |
| | DI | 1.22 | 0.44 |
| | NAc-c | 1.04 | 0.41 |
| | CPu | 1.04 | 0.33 |
| | PIR3 | 1.01 | 0.37 |
| | AI-v | 0.92 | 0.33 |
| | VSR-u | 0.92 | 0.49 |
| | GI | 0.88 | 0.52 |
| | VP | 0.86 | 0.38 |
| | Endo | 0.80 | 0.29 |
| | V-MID | 0.72 | 0.29 |
| | Am-u | 0.71 | 0.37 |
| | NAc-s | 0.63 | 0.29 |
| | AI-p | 0.62 | 0.43 |
| | PIR1 | 0.51 | 0.46 |
| | CLA | 0.47 | 0.30 |
| | Fr3 | 0.32 | 0.59 |
| | BNST | 0.25 | 0.37 |
| | M1 | 0.08 | 0.54 |
| | AI-d | -0.12 | 0.54 |
| | M2 | -0.23 | 0.59 |
| | IL | -0.31 | 0.27 |
| | LO | -0.31 | 0.46 |
| | S1 | -0.39 | 0.67 |
| | PrL | -0.47 | 0.43 |
| | OB | -0.53 | 0.52 |
| | Cg1 | -0.82 | 0.51 |
| | Sep | -0.86 | 0.60 |
| | PER | -1.26 | 0.58 |
| | LEC | -1.39 | 0.69 |
| | S2 | -1.50 | 0.54 |
| | VO | -1.60 | 0.55 |
| | PIR2 | -2.20 | 0.67 |
| | DLO | -2.24 | 0.75 |

| | | | | |
|---|---|---|---|---|
| | C | CA1 | 2.11 | 0.82 |
| | | V2M | 1.40 | 0.66 |
| | | GPe-m | 0.72 | 0.84 |
| | | ZI-A13 | 0.63 | 0.99 |
| | | STh | 0.04 | 0.64 |
| | | ZI-A11 | 0.00 | 0.74 |
| | | Cg2 | -0.09 | 0.64 |
| | | HTh-u | -0.30 | 0.63 |
| | | ZI | -0.52 | 0.70 |
| | | lPPC | -0.56 | 0.94 |
| | | GPe-l | -1.09 | 0.93 |
| | | PV | -1.14 | 0.54 |
| | | MO | -1.22 | 0.94 |
| Exercise | | | | |
| | A | VP | 1.10 | 0.69 |
| | | VSR-u | 1.10 | 0.65 |
| | | PrG | -0.73 | 0.76 |
| | | SMn | -0.73 | 0.72 |
| | | MO | -0.73 | 0.48 |
| | B | ANT | 1.43 | 0.75 |
| | | ZI | 0.33 | 0.87 |
| | | EP | 0.15 | 0.87 |
| | | STh | -0.95 | 0.60 |
| | | MHb | -0.95 | 0.50 |
| | C | Au1 | 1.07 | 0.70 |
| | | DG | 0.91 | 0.60 |
| | | DLO | 0.74 | 0.70 |
| | | Am-u | 0.74 | 0.72 |
| | | CA3 | 0.43 | 0.61 |
| | | AI-p | 0.42 | 0.67 |
| | | lPPC | -0.25 | 0.77 |
| | | mPPC | -0.37 | 0.77 |
| | | PER | -0.56 | 0.77 |
| | | CA2 | -0.81 | 0.74 |
| | | FC | -2.33 | 0.57 |
| | D | Cg2 | 1.56 | 0.64 |
| | | PrL | 1.28 | 0.61 |
| | | BFR-u | 1.12 | 0.66 |
| | | VO | 1.00 | 0.72 |
| | | V-MID | 0.19 | 0.70 |
| | | GPe-l | 0.10 | 0.59 |
| | | LO | 0.07 | 0.57 |
| | | VLO | -0.31 | 0.72 |
| | | BNST | -0.41 | 0.62 |
| | | OB | -1.02 | 0.69 |
| | | LEC | -1.03 | 0.69 |
| | | LD | -1.26 | 0.75 |
| | | LP | -1.29 | 0.75 |

| | | | |
|---|---|---|---|
| E | VENT | 0.97 | 0.80 |
| | Au2 | 0.64 | 0.78 |
| | RS | 0.55 | 0.84 |
| | CA1 | -1.07 | 0.88 |
| | S2 | -1.09 | 0.71 |
| F | Endo | 1.29 | 0.42 |
| | MD | 1.07 | 0.46 |
| | AI-v | 1.06 | 0.54 |
| | IMD | 0.99 | 0.52 |
| | Sep | 0.92 | 0.44 |
| | PIR3 | 0.89 | 0.49 |
| | Cg1 | 0.83 | 0.51 |
| | CLA | 0.76 | 0.60 |
| | PIR2 | 0.73 | 0.44 |
| | ZI-A11 | 0.65 | 0.56 |
| | CPu | 0.65 | 0.54 |
| | RT | 0.56 | 0.52 |
| | DI | 0.43 | 0.64 |
| | M1 | 0.41 | 0.60 |
| | AI-d | 0.39 | 0.66 |
| | M2 | 0.39 | 0.56 |
| | GPe-m | 0.32 | 0.30 |
| | NAc-c | 0.23 | 0.54 |
| | ILM | 0.20 | 0.36 |
| | PT | 0.18 | 0.14 |
| | S1 | 0.11 | 0.69 |
| | GI | 0.04 | 0.67 |
| | IL | -0.19 | 0.56 |
| | NAc-s | -0.43 | 0.59 |
| | Po | -0.49 | 0.69 |
| | LHb | -0.69 | 0.53 |
| | Fr3 | -1.03 | 0.64 |
| | ZI-A13 | -1.05 | 0.10 |
| | PIR1 | -1.07 | 0.71 |
| | PV | -1.63 | 0.12 |
| | V2M | -1.79 | 0.40 |
| | HTh-u | -1.95 | 0.00 |
| | IGL | -2.78 | 0.75 |
